# Phenotypic and metabolic plasticity shapes life-history strategies under combinations of abiotic stresses

**DOI:** 10.1101/328062

**Authors:** Lidor Shaar-Moshe, Ruchama Hayouka, Ute Roessner, Zvi Peleg

**Author notes:** **Corresponding author:** Zvi Peleg, The Robert H. Smith Institute of Plant Sciences and Genetics in Agriculture, The Hebrew University of Jerusalem. P.O. Box 12, Rehovot 7610001, Israel.

## Abstract

Plants developed various reversible and non-reversible acclimation mechanisms to cope with the multifaceted nature of abiotic stress combinations. We hypothesized that in order to endure these stress combinations, plants elicit distinctive acclimation strategies through specific trade-offs between reproduction and defense. To investigate *Brachypodium distachyon* acclimation strategies to combinations of salinity, drought and heat, we applied a system biology approach, integrating physiological, metabolic and transcriptional analyses. We analyzed the trade-offs among functional and performance traits, and their effects on plant fitness. A combination of drought and heat resulted in escape strategy, while under a combination of salinity and heat, plants exhibited avoidance strategy. On the other hand, under combinations of salinity and drought, with or without heat stress, plant fitness (i.e. germination rate of subsequent generation) was severely impaired. These results indicate that under combined stresses, plants’ life-history strategies were shaped by the limits of phenotypic and metabolic plasticity and the trade-offs between traits, thereby giving raise to distinct acclimations. Our findings provide a mechanistic understanding of plant acclimations to combinations of abiotic stresses and shed light on the different life-history strategies that can contribute to grass fitness and possibly to their dispersion under changing environments.

## Introduction

Phenotypic plasticity, the ability of a given genotype to express different phenotypes under alternative environmental conditions (Pigliucci, 2001), is a fundamental feature of plants, enabling them to optimize their fitness under changing and uncertain environments (Matesanz et al., 2010). The capacity of an organism to express plasticity in a given trait is mediated at the molecular level (Schlichting and Smith, 2002) and involves various reversible and non-reversible developmental, anatomical, morphological and physiological modifications (Nilson and Assmann, 2010). In contrast to genetic alterations that occur across several generations, phenotypic plasticity accounts for the plant responses to environmental changes within the life span of an individual plant (Bradshaw, 1965). It is assumed that the ability of phenotypic plasticity to alleviate short-term environmental changes could also be beneficial upon the long-term effects of climatic changes (Jump and Peñuelas, 2005).

Plant life-history, which aims at understanding and predicting the existence of plants with different traits and strategies, is predicted to change under environmental stresses through alterations in plant growth, reproduction and survival, the three main life-history traits (Grime, 1977). These performance traits, which directly contributing to plant fitness, can be modulated by morphological, physiological or phenological traits (i.e. functional traits) (Violle et al., 2007). Plant life-history strategies rely on trade-offs in resource partitioning between growth, reproduction and defense, due to resource limitations, and/or environmental stresses (Bazzaz et al., 1987). These constrains limit the capacity of the plants to acquire net carbon and alter the energy balance and allocation of assimilates from growth and reproduction into defense (Sultan, 2000). Variations in life-history strategies usually involve cross-species comparisons (e.g. (Salguero-Gomez et al., 2016). However, owing to their phenotypic plasticity, plants from the same genotype growing in different environments, can differ in the trade-offs of resource allocation, thus, exhibiting different life-history strategies (De Deyn, 2017). One of the challenges of plant biology is to identify these genetic trade-offs and through a mechanistic understanding manipulate them for crop improvement (Des Marais and Juenger, 2016).

Environmental stresses, such as soil salinization, drought episodes and heat waves, are among the main factors limiting plant performance and affecting their ecological fitness. At the Mediterranean-like conditions, these stresses usually occur simultaneously, effecting both natural and agricultural systems (Vautard et al., 2007; Maggio et al., 2011). Moreover, based on climate change models, plants are projected to encounter a greater range and number of abiotic stress combinations in the near future. To cope with such environmental constrains, plants have evolved a variety of acclimation mechanisms. These mechanisms can be encapsulated to two main strategies: *escape* (e.g. rapid completion of the plant life cycle or deeper rooting), and *resistance*, which is sub-divided to *avoidance* (altering or halting the effect of the stress through mainly morphological and/or anatomical changes), and *tolerance* (maintaining normal functions despite the presence of the stress through molecular mechanisms) (Levitt, 1972; Mickelbart et al., 2015). Plants can combine acclimation mechanisms from different strategies, thus eliciting an environmentally induced phenotypic variation.

*Brachypodium distachyon* (L.) Beauv. is a model system for Pooid cereal and forage species that has been used to study environmental stress responses such as: drought, extreme temperature, salinity and nutrient availability (reviewed by Scholthof et al., 2018). However, except for a few studies (e.g. Barhoumi et al., 2010; Des Marais et al., 2017; Shaar-Moshe et al., 2017), most of these experiments involved single stresses at early vegetative stages. In order to study *B. distachyon* acclimation strategies to environmental stresses, information on life-history and phenological data is required since they determine the coordination of environmental conditions with developmental transitions, the occurrence of the stress and the expected acclimation strategy (Des Marais and Juenger, 2016).

Recently we have shown a unique transcriptional signature among genes that were differentially expressed only under combinations of stresses. This transcriptional pattern was enriched with antagonistic responses, suggesting alterations in the mode of action under different combinations of stresses (Shaar-Moshe et al., 2017). Here we used a system biology approach to test the hypothesis that plants elicit a unique acclimation strategy, with distinct trade-offs in life-history traits of growth, reproduction and defense, under each combination. The objectives of this study were to ***i***) characterize the effects of combined stresses on plant fitness and ***ii***) determine the trade-offs among functional and performance traits. Our results shed light on life-history strategies, and the capacity of phenotypic and metabolic plasticity, which can facilitate plant acclimations to the projected changing environments.

## Material and methods

### Plant material and growth conditions

*Brachypodium distachyon* accession *Bd*21-3 plants were grown as previously described (Shaar-Moshe et al., 2017). Briefly, following 48h at 4°C to synchronize germination and 5d at 15°C to establish germination, uniform seedlings were transplanted into pre-weighted 1L pots and transfer to a Phytotron (22°C day/16°C night, 10h light/14h dark). The dry and fully irrigated weight of the pots were used to calculate soil water content. Plants were irrigated to runoff with fresh water three times a week, and fertilized with 1g L^−1^ N:P:K (20% nitrogen, 20% phosphorus and 20% potassium) + micronutrients, eight weeks after germination. Ten weeks after germination, plants transferred to a long day regime (15h light/9h dark).

Each plant was subjected to one of the following five treatments in a split-plot factorial (treatment) complete random design: *Control* (***C***), plants were grown at 22°C day/16°C night throughout the experiment. Combination of salinity and heat (***S_&_H***), plants were exposed progressively to salinity, starting at five-leaf stage, by two irrigations of 20mM NaCl, followed by five irrigations with 50 and then 80mM NaCl. Target concentration of 100mM NaCl, which was achieved within four weeks, was kept throughout the experiment. Runoff electric conductivity was monitored weekly. At anthesis, plants were transferred on 20:00 pm, to a pre-heated greenhouse at the Phytotron (34°C day/28°C night) for four days. Combination of drought and heat (***D_&_H***), plants were gradually exposed to water stress, starting at booting stage (BBCH scale 45; Hong et al., 2011, approximately 12 weeks after germination), by withholding irrigation. Each pot was weighted every second day and relative soil water content was maintained at 40% for 17 days. At anthesis, plants were subjected to heat stress as described above. Combinations of Salinity and drought (***S_&_D***) and (***S_&_D_&_H***), salinity and drought stresses were applied gradually at five-leaf and booting stage, respectively, as described above. Heat stress was applied at anthesis, as described above. Under the two combinations, plants were irrigated with fresh water once drought stress was imposed (Supplementary Fig. S1).

### Performance traits and plant fitness

Plants were documented and measured for shoot dry weight (DW) at three developmental stages during the reproductive stage: 4, 12 and 17 days after anthesis (DAA). At the end of the growing season, when plants from all treatments senesced, spikes and shoots, from each plant, were separately harvested (*n*=5) and oven-dried (75°C for 96h). Samples were weighted to determine shoot DW. The reproductive samples were threshed, and grain were weighted and counted to obtain 100-grain weight. The ratio between spike DW and whole shoot DW was used to calculate the reproductive allocation. Grain parameters were analyzed using GrainScan software (Whan et al., 2014). Germination assay was performed using the “cigar roll” method (Watt et al., 2013). Twenty uniform grains, which developed on plants subjected to control or stress combination treatments (*n*=3), were placed on a moist germination paper (25×38cm; Anchor Paper Co., St. Paul, MN, USA), three cm from the top of the sheet, with the embryo facing down. The paper was covered with additional sheet of moist germination paper and then rolled tightly to a final diameter of ~4 cm. The base of the rolls was dipped in a tray of tap water and stored in darkness for 48h at 4°C, followed by 6d at 15°C for establishment of germination. The number of grains with the primary seminal root was recorded to determine grain viability.

### Functional traits

Measurements of chlorophyll fluorescence and content were conducted in a complete random design at three developmental stages during the reproductive stage: 4, 12 and 17 DAA, on the mid portion of the adaxial side of the third leaf. Chlorophyll fluorescence was measured using a portable chlorophyll fluorometer MINI-PAM-II (Walz GmbH, Effeltrich, Germany). Dark-adapted *Fv*/*Fm* was determined following 40 min of whole plant dark-adaptation (*n*=4). *Fv*/*Fm* values that represent maximal photochemical efficiency of photosystem II were automatically calculated as (*Fm-F_0_*)/*Fm* by the WinControl-3 software. Chlorophyll content (SPAD value; *n*=4) was assessed by the SPAD-502 chlorophyll meter (Minolta Camera Co., Japan). Each SPAD value is the mean of five technical replicates measured from five different leaves, per plant. Carotenoid content and plant senescence were estimated using none-distractive reflectance measurements, on the mid portion of the adaxial side of the third leaf, by the CI-710 Miniature Leaf Spectrometer (CID Bio-Science, USA), at 4 DAA. Integration time was 800 milliseconds, boxcar width was 10 points and each value represents an average of four scans. The following reflectance indices were used for carotenoid fluorescence: CRI_550_ = (1 / W_510_) - (1 / W_550_) (Gitelson et al., 2002) and plant senescence: PSRI = (W_680_ – W_500_) / W_750_) (Merzlyak et al.,1999), where W represents the wavelengths used to calculate the indices. Measurements of relative water content (RWC) and osmotic adjustment (OA) were performed on the third leaves at mid-day (*n*=5), as described previously (Shaar-Moshe et al., 2015). Ion concentrations (i.e. Cl^−^, Na^+^ and K^+^), under control and stress treatments, were evaluated based on (Yermiyahu et al., 2017). Oven-dried leaf samples (*n*=6) (75°C for 96h) were ground to a fine powder. One hundred mg from each sample was added to 10 mL double-distilled water (DDW), followed by an overnight incubation. Sodium (Na^+^) and potassium (K^+^) concentrations were analyzed by atomic absorption (PerkinElmer Precisely Analyst 200) and chloride (Cl^−^) concentrations were quantified with chloridometer (MK II Chloride Analyzer 926, Sherwood Scientific Ltd., Cambridge, UK).

### Transcriptional data and analysis

Extraction of RNA from flag and second leaf samples and preparation of cDNA libraries and their sequencing were previously described in Shaar-Moshe et al. (2017). Briefly, at 4 DAA leaf samples, were collected, frozen in liquid nitrogen and total RNA was extracted using Plant/Fungi Total RNA Purification Kit (Norgen Biotek Corp., Canada). Three biological replicates from each condition, with the highest values of RNA integrity, chosen for RNA sequencing. Twenty-four cDNA libraries were subjected to single-end multiplex sequencing (50bp) on the Illumina HiSeq2500 sequencer (Technion Genome Center, Haifa, Israel), using the TruSeq RNA sample preparation kit ver.2 (Illumina, San Diego, USA), according to manufacturer’s standard protocols. Data processing, which included barcode removal, filtering of low quality reads and alignment of the reads to the *B. distachyon* reference genome (Bd21-3 v.1), as well as analysis of differentially expressed genes, with DESeq2 package (Love et al., 2014), and their functional annotation, based on MapMan tool (Thimm et al., 2004), were obtained form (Shaar-Moshe et al., 2017). Files of the raw mRNA-seq are available at the short-read archive of the National Center for Biotechnology Information (https://www.ncbi.nlm.nih.gov/sra) under accession number PRJNA360513.

### Determination of starch content

Starch content was determined as described previously (Naschitz et al., 2010), with minor modifications. Briefly, 50mg of lyophilized leaf samples (*n*=5) that were collected at 5 DAA at 18:00 pm were ground with liquid nitrogen and rinsed three times with 80% ethanol to remove soluble sugars. The pellet was fluidized with DDW and starch was hydrolyzed with amyloglucosidase (Sigma), in 15mM sodium acetate pH 4.5, overnight at 55°C. Subsequently, soluble sugar content was determined spectrophotometrically with Epoch plate reader, with absorbance at 340nm, using Glucose (HK) assay kit (Sigma).

### Metabolite extraction

Flag and second leaf (*n*=5) were harvested at four DAA, immediately frozen in liquid nitrogen and stored at −80°C until lyophilization. Sample were weighted and transferred to cryo-mill tubes (Precellys 24, Bertin Technologies). Methanol (MeOH, 500 μL) containing the internal standards [D-Sorbitol-^13^C_6_ (0.02 mg/mL) and L-Valine-^13^C_5_,^15^N (0.02mg/mL), Sigma Aldrich (Australia)] was added to the samples, followed by vortex and homogenization (3×45s at 6400 rpm) at −10°C using a Cryomill (Precellys 24, Bertin Technologies). The samples were then extracted at 70°C in a thermomixer at 850 rpm, and centrifuged for 5 min at 4°C, at 13,000 rpm. The MeOH supernatant was collected into new reaction tubes and 500μL water (Milli Q grade) was added to the tubes containing the sample pellet, followed by vortex-mixed for 30s, and centrifuged at 13,000 rpm for 10 min at 4°C. The supernatants were combined and 100μL were dried *in vacuo* for gas chromatography–mass spectrometry (GC–MS) untargeted analysis.

### Derivatization for GC–MS analysis

Derivatization for GC–MS analysis was performed as described in Dias *et al.* (2015). After completion of derivatization, samples were set for one hour and then 1μL was injected onto the GC column using a hot needle technique. Splitless and split (1:20) injections were performed for each sample.

### Untargeted GC–MS and statistical analysis

Untargeted GC–MS analysis and data analysis were carried out as described in (Hill et al., 2015). The metabolite data were analyzed with the open-source software, MetaboAnalyst 3.0 (Xia and Wishart, 2016). Interquartile range was used for data filtering, followed by log transformation for data normalization. The metabolites are presented as fold change (FC) values relative to control conditions and a 10% false discovery rate correction for multiple comparisons (Benjamini and Hochberg, 1995), was used to detect significant alterations in metabolite accumulation.

### Statistical and data analyses

Measurements of functional and performances traits, as well as correlation between gene expression data and functional traits or metabolites were analyzed statistically using JMP^®^ pro 13 statistical package (SAS Institute, Cary, NC, USA). All traits were examined for homoscedasticity among treatments. Differences between control and stress treatments, were detected using one-way analysis of variance (ANOVA), followed by Dunnet’s test at *P*≤0.05. In the absence of homoscedasticity among treatments, nonparametric comparisons with Steel method (*P*≤0.05) were used to detect differences between control and stress conditions. Differences between two treatments were detected using Student’s *t*-test at *P*≤0.05. Principle component analysis (PCA), based on log_2_ FC values of genes assigned to functional categories (e.g. carotenoid synthesis and starch metabolism), was used to identify the first principal component (PC1, eigenvalues >1) that accounted for the maximal variation among gene expression. Associations between PC1 and the FC values of the corresponding functional traits or metabolites were studied using Pearson correlation analysis at *P*≤0.1. In the lack of liner relationship, nonparametric correlation with Spearman’s correlation was used. Correlations between traits or between metabolites were studied using Pearson correlation analysis at *P*≤0.05. PCA was used to evaluate the association between osmolytes (i.e. 11 metabolites and 3 ions) and OA and RWC. PCA is presented as a biplot of the two major PCs (Eigenvalues >1). Line and box plots were generated by ggplot function in ggplot2 package.

## Results

### Under S_&_H plants maintained their reproductive allocation, while under S_&_D and S_&_D_&_H their fitness was severely impaired

All stress combinations resulted in detrimental phenotypic effects, which included a reduction in shoot height and chlorophyll content (SPAD units; Fig. 1A). However, depending on the combination of the stresses, a differential acclimation response, which varied in magnitude and timing, was elicited by the plants. For example, under the combination of drought and heat (D_&_H), chlorophyll content decreased by 20 and 82%, at 4 and 12 DAA (i.e. at grain filling), respectively, compared with control. Yet, under the other stress combinations, significant reductions in chlorophyll content were detected only at 17 DAA (during grain maturation period) (Fig. 1B, Supplementary Table S1). Genes involved in chlorophyll degradation were up-regulation at four DAA, among all combined stresses. A predominant increase was found under D_&_H that resulted in the highest expression of *chlorophyll b reductase* (BRADI1G13416), which catalyzes the first step in the conversion of chlorophyll b to a, and *pheophorbide a oxygenase* (BRADI3G53860 and BRADI1G75270), which produces red chlorophyll catabolite from pheophorbide a (Barry, 2009) (Supplementary Table S2).

**Fig. 1.**
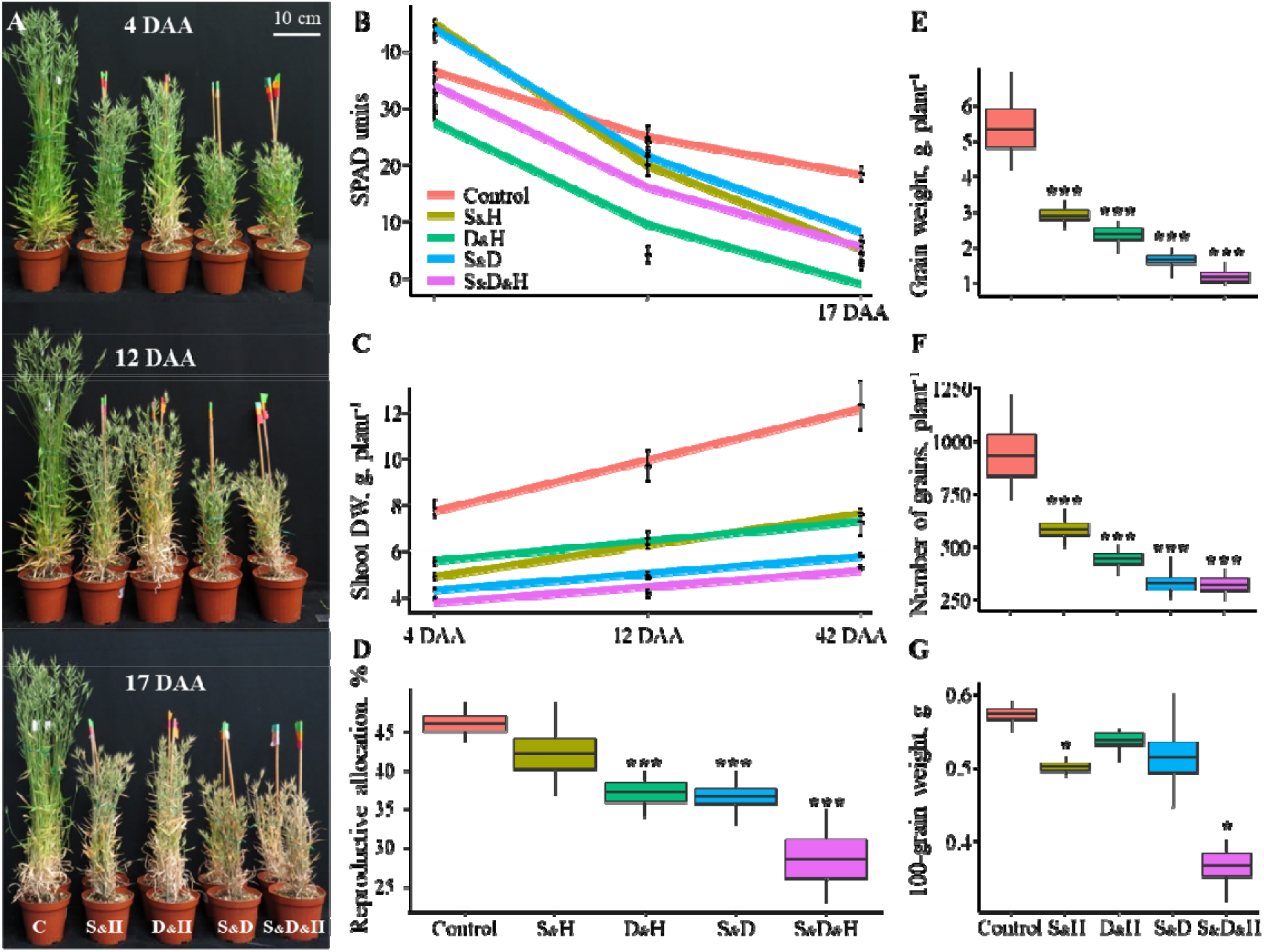
Performance traits and fitness of *Brachypodium distachyon* plants grown under control (C) and combinations of stresses: salinity and heat (S_&_H), drought and heat (D_&_H), salinity and drought (S_&_D), salinity, drought and heat (S_&_D_&_H). (A) Plants grown under control conditions and four combinations of stresses at three developmental stages during the reproductive stage: 4, 12 and 17 days after anthesis (DAA). (B) Chlorophyll content at 4, 12, and 17 DAA, under control and combinations of stresses. (C) Accumulation pattern of shoot dry weight (DW) at 4, 12, and 42 DAA. Values and significance levels are indicated in Supplementary Table S1. (D) Box plots of reproductive allocation (i.e. the ratio between spike and shoot DW), (E) grain weight, (F) number of grains per plant, and (G) 100-grain weight. * and *** indicate significant differences between control and stress treatments at *P*≤0.05 and 0.001, respectively, as determined by Dunnet’s test. Values are mean (*n*=4)±SE.

To assess the effects of the combined stresses on plant fitness and its performance traits, we measured shoot dry weight (DW) at three reproductive stages; 4, 12 and 42 DAA (i.e. plant senescence). In addition, we measured spike DW at 42 DAA and estimated the proportion of spike DW per shoot DW, as a surrogate for reproductive allocation. Overall, the plants accumulated significantly less shoot DW under the combined stresses, compared with control conditions. Under D_&_H and the combination of salinity and heat (S_&_H), plants accumulated on average 34% more DW than under the combination of salinity and drought (S_&_D) and salinity, drought and heat (S_&_D_&_H), across the three developmental stages. During the reproductive stages the vegetative growth of cereals is ceased, thus, any accumulation of shoot DW is attributed to an increase in the weight of reproductive tissues and subsequently to grain production. Under control and S_&_H, the plants accumulated on average ~55% of their weight from 4 DAA until senescence (i.e. 42 DAA) and showed a comparable level of reproductive allocation. However, the other stress combinations showed a milder DW accumulation of only ~33% and their reproductive allocation was significantly decreased, especially under S_&_D_&_H (Fig. 1C, D). To assess the causes for the different spike DW and reproduction allocation among the combined stresses, we examined several yield parameters. Total grain weight and number followed a similar gradual decrease as shoot DW. Under S_&_H, plants showed limited grain loses, while under S_&_D_&_H plants exhibited sever impairments (Fig. 1E, F). This pattern was not observed upon examination of 100-grain weight, which showed significant decreases under S_&_H and S_&_D_&_H, compared with control (Fig. 1G). As shoot DW accumulated to similar values under S_&_H and D_&_H, the difference in the reproductive allocation between the two combinations, can be attributed to higher values of total grain weight and number, under S_&_H, which compensated for the mild decrease in 100-grain weight (Fig. 1C-G). Examination of grain viability, of grains that developed on plants subjected to combinations of stresses, revealed a severe impairment in germination rate only under S_&_D and S_&_D_&_H, compared with control (Supplementary Fig. S2).

### D_&_H resulted in a rapid reduction of chlorophyll florescence

To investigate the effects of the combined stresses on the photosynthetic apparatus and its associated pigments, we measured the maximal photochemical efficiency of photosystem II (PSII) (*Fv*/*Fm*) at the three developmental stages during the reproductive stage. Under both S_&_D_&_H and D_&_H, *Fv*/*Fm* declined by ~12 and 92%, at 12 and 17 DAA, respectively, compared with control (Fig. 2A and Supplementary Table S1). Physiological responses and acclimation are the consequences of several level of regulations, including orchestrated transcriptional changes. To evaluate the contribution of transcriptional modifications to plant acclimation under combined stresses and identify potential pathways that may directly regulate plant phenotypes, we examined the association between physiological traits and the transcriptional patterns of the underlying genes. This analysis revealed a high correlation (r=0.9, *P*=0.1) between the physiological changes of PSII efficiency and expression patterns of genes that are involved in light harvesting complex and polypeptide subunits of PSII (Fig. 2A, B and Supplementary Table S2). Genes that showed a higher expression level under S_&_H and S_&_D include *PSB27-H1* (BRADI1G63300), which is required for the formation and stability of PSII-LHCII supercomplexes in *Arabidopsis*, and *PSB27-H2/LPA19* (BRADI1G08720), which functions in C-terminal processing of D1. Among the genes that displayed a lower expression level under S_&_D_&_H and D_&_H is *PPL1* (BRADI1G77047) that is required for efficient repair of photodamaged PSII (Lu, 2016).

**Fig. 2.**
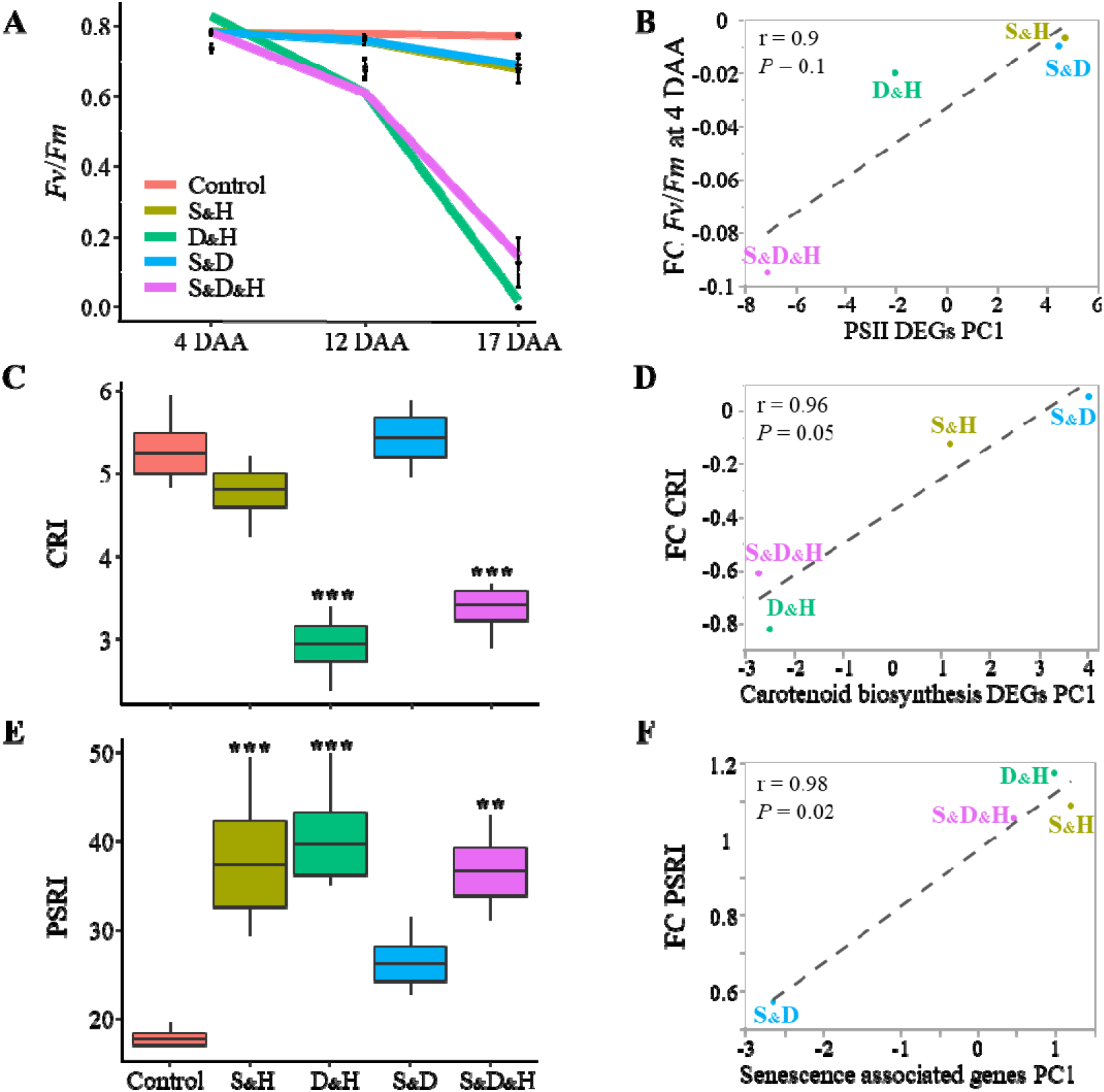
Differential effects of combined stresses on photosynthetic apparatus, its associated pigments and the progression of senescence at the physiological and transcriptional levels. (A) Maximal photochemical efficiency of photosystem II (*Fv*/*Fm*), at 4, 12, and 17 days after anthesis (DAA), under control and combinations of stresses. (B) Correlation between first principle component (PC1) of expression pattern of genes related to photosystem II (PSII) and fold change (FC) values of *Fv*/*Fm* at 4 DAA. (C) Box plots of carotenoid reflectance index (CRI) that is calculated as 1/W510 - 1/W550. W represents the wavelengths used to calculate the reflectance index. (D) Correlation between PC1 of expression pattern of genes involved in carotenoid synthesis and FC values of carotenoid content. (E) Box plots of plant senescence reflectance index (PSRI) that is calculated as [(W680-W500) / W750] × 1000. (F) Correlation between PC1 of expression pattern of senescence associated genes (SAGs) an PSRI. ** and *** indicate significant differences between control and stress treatments at *P*<0.01 and 0.001, respectively, as determined by Dunnet’s test. Values are mean (*n*=4)±SE. Growth conditions are as follows: salinity and heat (S_&_H), drought an heat (D_&_H), salinity and drought (S_&_D), salinity, drought and heat (S_&_D_&_H).

### S_&_H and S_&_D differentially affect carotenoid content and leaf senescence

Carotenoids have a structural role in the assembly of the photosynthetic apparatus, and the retention of carotenoids, in the progress of chlorophyll breakdown during senescence, has been suggested as a mechanism of photoprotection (Nisar et al., 2015). To assess the content of carotenoid in the leaves, we used the non-destructive carotenoid reflectance index 1 (CRI1) (Gitelson et al., 2002). Under D_&_H and S_&_D_&_H, plants exhibited low values of CRI1, whereas under S_&_H and S_&_D, CRI1 values were comparable to control (Fig. 2C). In accordance, the expression pattern of genes that are involved in carotenoid metabolism exhibited a high and positive association with carotenoid content under the combined stresses (Fig. 2D and Supplementary Table S2).

Plant senescence reflectance index (PSRI), which estimates the proportion between chlorophyll and carotenoids (Merzlyak et al., 1999), was used to quantitatively evaluate leaf senescence. All stress combinations, except for S_&_D, showed higher values of PSRI, compared with control, indicating stress induced-senescence (Fig. 2E). This pattern was also reflected at the transcriptional level by a strong association between PSRI and senescence associated genes (SAG). In addition, a high correlation (r=0.97, *P*=0.03) was detected between PSRI and genes involved in chloroplast nucleoid metabolism and maintenance (Supplementary Table S2), a pathway that was enriched among genes uniquely expressed only under stress combinations (Shaar-Moshe et al., 2017). The different PSRI values that were detected between S_&_H and S_&_D might be attributed to a higher carotenoid content under S_&_D, as chlorophyll contents were comparable under these stress combinations (Figs. 1B, 2C). Consistently, mRNA levels of *phytoene synthase* (*PSY*, BRADI4G37520), a rate-limiting enzyme, which catalyzes the first committed step in carotenogenesis (Hirschberg, 2001), were up-regulated only under S_&_D (Supplementary Table S2).

### Starch content depleted under heat-related stress combinations

Under abiotic stresses, such as salinity, drought and heat, starch is usually degraded to support the plant energy demands and carbon supply at times when photosynthesis is limited (Thalmann and Santelia, 2017). This process resulted in the accumulation of maltose, a major degrading product of starch, and of its deriving sugars (Thalmann et al., 2016). Under all heat stress combinations, plants accumulated less starch in their leaves, compared with control conditions (ranging between 65% and 97%; Fig. 3 and Supplementary Fig. S3A). The pattern of starch attenuation, correlated with expression pattern of starch synthesis and degradation DEGs (Supplementary Table S2 and Fig. S3B), suggesting a tight regulation, of both anabolic and catabolic processes, under the combined stresses. In addition, starch was in a close association with maltose content (r=1, *P*=0.004; Supplementary Fig. S3C). Starch content in the leaves did not correlate with photosynthesis rate (r=0.62, *P*=0.3; Supplementary Fig. S3D) or with grain weight (r=0.54, *P*=0.3, Supplementary Fig. S3E). The plasticity of starch attenuation under the different stress combinations demonstrates that it cannot be merely considered as a storage compound (Thalmann and Santelia, 2017) and may imply its involvement in distinct shifts between defense, maintenance and reproductive processes.

**Fig. 3.**
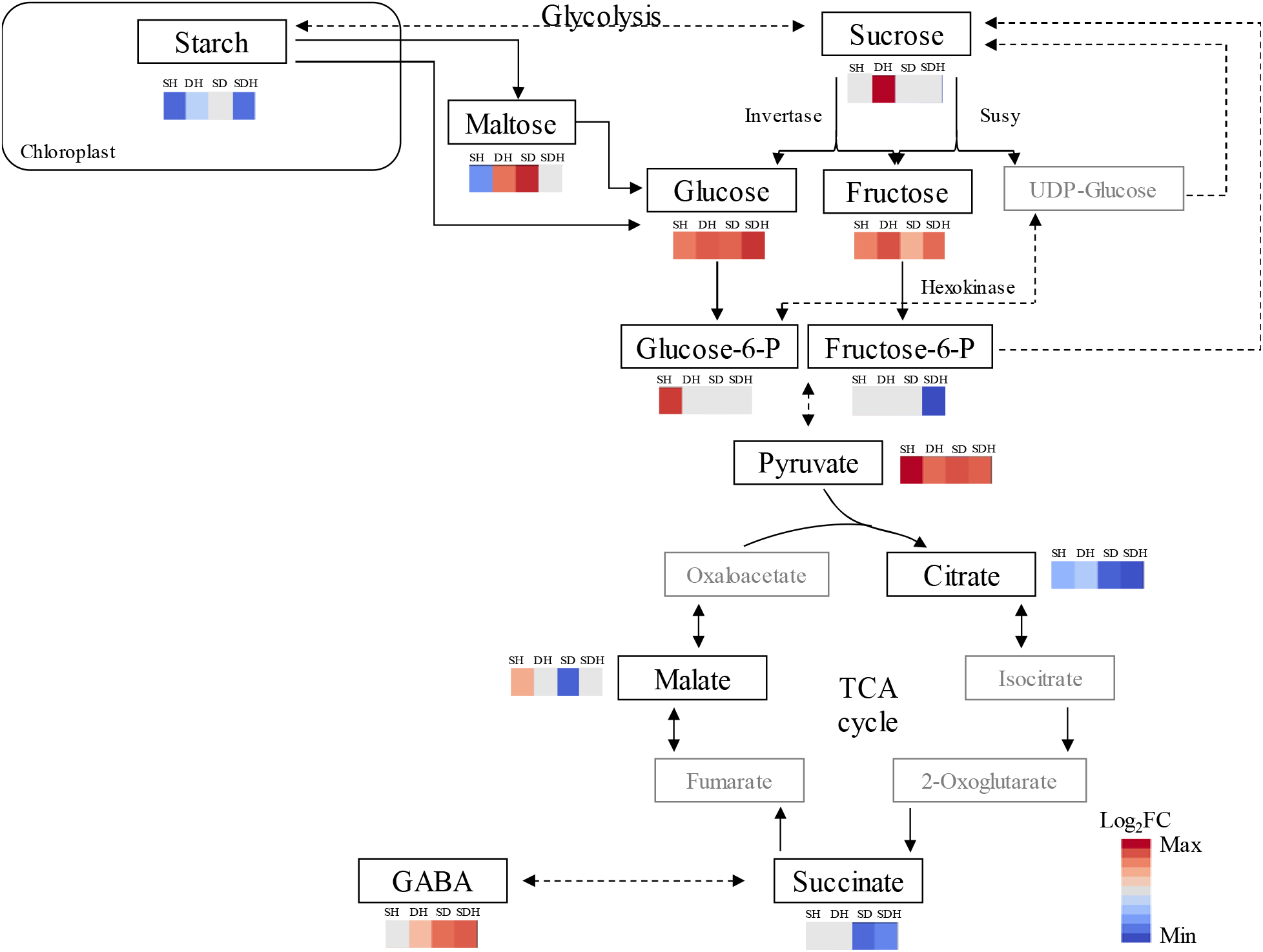
Accumulation patterns of metabolites involved in energy metabolism in *Brachypodium distachyon* plants grown under control and combinations of stresses: salinity and heat (S_&_H), drought and heat (D_&_H), salinity and drought (S_&_D), salinity, drought and heat (S_&_D_&_H). Red and blue colors indicate significant (*P*≤0.1) accumulation and depletion of metabolite content, respectively, and white color indicates a non-significant change, compared with control.

### Plant energy balance is severely impaired under S_&_D and S_&_D_&_H

Starch degradation and remobilization under stress releases sugars and other derived metabolites, which can support plant growth, function as compatible solutes and provide an alternative source of energy and carbon (Krasensky and Jonak, 2012). To improve our understanding on cellular energy metabolism under stress, we performed a complete foliar metabolic profiling, in parallel with the transcriptional analysis. Among the 70 annotated metabolites, differentially accumulated under at least one stress combination, most metabolites (94%) showed the same response pattern (i.e. accumulation or depletion), across the four combinations. However, the intensity of the response varied among the different combinations (e.g. GABA, tryptophan and mannose reached FC differences of 71, 48, and 22, respectively, compared with control conditions; Supplementary Table S3). Functional classification of the metabolic data revealed that sugars, sugar phosphates and sugar alcohols constituted up to 41% of the metabolites and enrichment analysis further identified galactose metabolism as a prevalent pathway under stress. This pathway includes sugars with various functions such as signaling (e.g. glucose and fructose), transport (sucrose and raffinose), osmoprotectants (e.g. galactinol and myo-inositol), cell wall polysaccharides (e.g. galactose and mannose) and glycerolipid metabolism (e.g. galactosylglycerol), which showed distinct metabolic responses under each stress combination (Supplementary Fig. S4).

We further focused our analysis on metabolites that are involved in cellular respiration. Sucrose and its two hexoses constitute the early steps of glycolysis, the first stage of respiration. Plants accumulated sucrose only under D_&_H (FC=1.2). While its breakdown products, the hexoses fructose and glucose, accumulated under all combined stresses (Fig. 3 and Supplementary Table S3). The hexoses are further converted to glucose-6-P that accumulated under S_&_H (FC=2.2) and fructose-6-P, which depleted under S_&_D_&_H (FC=-4.6). The endproducts of glycolysis include the organic acid pyruvate that accumulated to comparable levels under all stress combinations and malate, which accumulated under S_&_H and decreased under S_&_D (FC=1.8 and −3.4, respectively). Additional organic acids of the tricarboxylic acid (TCA) cycle, showed reduced levels under the combined stresses. For example, succinate, a metabolite connecting the TCA cycle and γ-Aminobutyric acid (GABA) shunt (Fait et al., 2007), showed decreased levels only under S_&_D and S_&_D_&_H. Consistently, high levels of GABA were detected in leaves of plants subjected to these two stresses (Fig. 3 and Supplementary Table S4).

### Preferable accumulation of osmolytes under combined stresses

Accumulation of compatible solutes is an adaptive mechanism that mitigates the deleterious effects of an osmotic stress by sustaining cellular turgor and water status and protecting cellular constituents (Rontein et al., 2002). Under all combined stresses, an increase in osmotic adjustment (OA) was detected, predominantly under S_&_D and S_&_D_&_H. However, this increment was not sufficient to retain leaf relative water content (RWC), to a level comparable to control conditions. Heat stress did not contribute to the OA, as indicated by the similar values of OA under S_&_D and S_&_D_&_H, but it led to a dramatic loss of 20% of RWC under S_&_D_&_H compared with S_&_D (Fig. 4A).

**Fig. 4.**
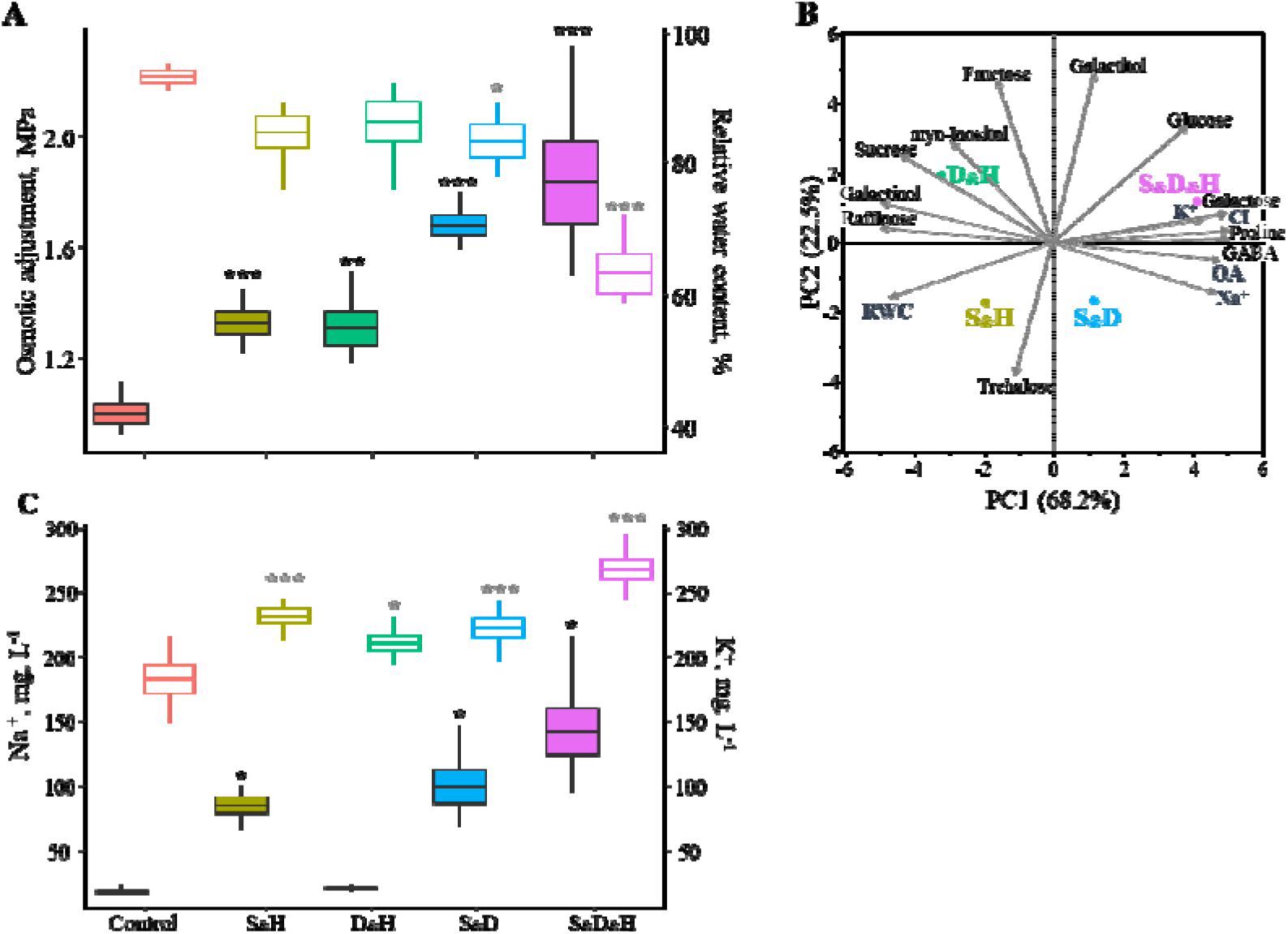
The contribution of osmolytes to osmotic adjustment (OA) and its relationship with relative water content (RWC). (A) Box plot of OA and leaf RWC, denoted by filled and empty boxes, respectively. (B) Biplot of a principal component (PC) analysis of the relationship among osmolytes (i.e. 11 organic osmolytes, detected by metabolic profiling, and three inorganic osmolyes) and the physiological traits OA and RWC, under combinations of stresses. (C) Box plot of Na^+^ and K^+^ concentrations, denoted by filled and empty boxes, respectively. *, ** and *** indicate significant differences between control and stress treatments at *P*≤0.05, 0.01 and 0.001, respectively, as determined by Dunnet’s test. Values are mean (*n*=6)±SE. Growth conditions are as follows: salinity and heat (S_&_H), drought and heat (D_&_H), salinity and drought (S_&_D), salinity, drought and heat (S_&_D_&_H).

To gain a comprehensive understanding of the relationship between osmolytes (organic and inorganic) and OA and their effect on RWC, we performed a principle component analysis (PCA) that included 11 metabolites, three ions and the two physiological measurements. The cumulative contribution of the two major principal components (PCs, Eigenvalues >1), to the total variation, was 91%. PC1 that explained 68.2% of the variation was loaded positively with OA, the metabolites proline, GABA, galactose and glucose, and the ions Na^+^, K^+^ and Cl^−^, suggesting the involvement of these osmolytes in osmoregulation. On the other hand, PC1 was loaded negatively with RWC, raffinose, galactinol, sucrose and myo-Inositol, indicating the negative association between OA and RWC and implying that these metabolites did not contribute to osmotic adjustment in *B. distachyon* (Fig. 4B). The lack of correlation between OA and fructose, galactitol and trehalose may result from additional functions these metabolites hold, such as signaling and transport (Ruan, 2014). Cumulative FC of significant compatible solutes revealed their gradual accumulation across the combined stresses, ranging between 39, under S_&_H, and reaching up to 129 cumulative FC, under S_&_D_&_H (Supplementary Table S3). Although S_&_H and D_&_H had comparable levels of OA (Fig. 4A), under D_&_H plants accumulated a higher level of compatible solutes compared with S_&_H (i.e. cumulative FC of 50 *versus* 39, respectively).

Salt-treated plants accumulated higher levels of Na^+^ in their leaves compared with control and D_&_H. Accumulation of K^+^, which was positively correlated with both Na^+^ and Cl^−^ accumulation, was detected under all combined stresses, especially in plants subjected to S_&_D_&_H that accumulated 45% more K^+^, compared with control (Fig. 4C). The accumulation patterns of K^+^ and Na^+^ across the combined stresses resulted in a similar four-fold reduction in K^+^ / Na^+^ ratio among the salt-treated plants (Supplementary Fig. S5A). In addition, Cl^−^ was accumulated under all combined stresses, primarily in plants subjected to S_&_D_&_H, which accumulated twice the amount of Cl^−^, compared with control conditions (Supplementary Fig. S5B).

## Discussion

Plant acclimations to abiotic stress conditions occur through a coordinated sequence of transcriptional, translational, metabolic and physiological events, from the cellular to the whole plant level. These processes change the metabolic status of the plants toward a new steady-state level that supports plant growth and fitness under the unfavorable conditions. To study acclimation mechanisms of temperate grasses to combinations of abiotic stresses, we subjected *B. distachyon* plants to combinations of salinity, drought and heat stresses, which are the major environmental constrains at the Mediterranean region (Vautard et al., 2007;Maggio et al., 2011). We examined the extent of phenotypic and metabolic plasticity and the limitation such conditions imposed on plant performances and fitness. Our analysis revealed commonalities and divergences between the life-history strategies plants elicited under each unique stress combination.

### Under D_&_H plants exhibited a life-history strategy of accelerated senescence

The Mediterranean-like environments are usually affected by prolonged water stress combined with heat waves, causing together extensive agricultural losses (Loss and Siddique, 1994). Under such conditions, annual plants that have a rapid life cycle and escape the spring heat waves may gain a high fitness. The *B. distachyon* accession Bd21-3 that was collected in northern Iraq is native to an area with low winter precipitation and high summer temperatures (Des Marais and Juenger, 2016). Accordingly, these plants exhibited an accelerated senescence under D_&_H that was manifested by a significant reduction in chlorophyll content, starting as early as 4 DAA (Fig. 1B), a reduction in photosynthesis rate (Shaar-Moshe et al., 2017), accumulation of galactosylglycerol, consistent with breakdown of thylakoid membranes, which contain up to 80% galactosylglycerolipids (Holzl and Dormann, 2007), and accumulation of transport sugars (e.g. sucrose and galactitol) (Fig. 4 and Supplementary Fig. S4 and Table S3). Leaf senescence, which is characterized by the transition from nutrient assimilation to nutrient remobilization, involves, in its early phase, breakdown of chloroplasts and decrease in photosynthesis (Avila-Ospina et al., 2014). The positive correlation between leaf senescence (measured by PSRI) and expression pattern of genes that are involved in chloroplast nucleoid metabolism and maintenance provides a transcriptional support for the link between leaf aging and chloroplast stability. The associations between physiological traits and their underlying gene expression patterns suggest a coordinated regulation between the transcriptional and physiological level and may facilitate detection of pathways and genes that directly control plant acclimations to stress conditions.

Despite a reduction in photosynthesis under D_&_H, the hexoses, glucose and fructose accumulated under D_&_H (Fig. 3 and Supplementary Table S3). This may occur through starch remobilization, which provides an alternative source of energy and carbon, or through a decrease in amino acid synthesis, which leads to a higher carbon availability (Jongebloed et al., 2004). Indeed, under D_&_H, starch level depleted, compared with control conditions, and the amino acids tryptophan and GABA, only mildly accumulated, compared with plants grown under the other stress combinations (Supplementary Table S3). In addition, considering that up to 70% of the leaf proteins are localized within the chloroplasts, the recycling and management of materials accumulated during leaf development into exportable nutrients, is a critical process in senescing plants to sustain yield and fitness (Sade et al., 2018). The accumulation of transport sugars that can move through the phloem to the filling grains may indicate that nutrient remobilization was accelerated under D_&_H. Yet, nutrient reservoir and/ or their transport rate did not meet the progression of senescence, resulting in a reduction of 19% in reproductive allocation and a decreased in total grain weight (Fig. 1D-E). This life-history strategy of accelerated senescence resulted in a trade-off in the reproductive allocation via shortened grain-filling period, which consequently diminished grain yield.

### Plants grown under S_&_H exhibited moderate effects of the stress on functional and performance traits

Plants subjected to S_&_H were able to sustain, for a longer period, higher or comparable values of chlorophyll content and *Fv*/*Fm*, respectively (Figs. 1B, 2A), which further support the moderate decreases in photosynthesis rate, compared with control (Shaar-Moshe et al., 2017). Under S_&_H plants accumulated 29% less shoot DW, similarly to D_&_H, yet the reproductive allocation was comparable to control conditions, indicating that the plants maintained source-sink relationships and successfully partitioned photosynthetic assimilates to the reproductive tissues (Fig. 1C, D). Plants grown under S_&_H also maintained leaf water content at a level comparable to control conditions, possibly through an increase in OA (Fig. 4A). By increasing the intercellular osmolarity plants maintain a high cellular turgor that is necessary for cell expansion and facilitate a higher stomatal conductance under lower water potentials, thus actively modifying the environment they experience.

Osmotic adjustment can be achieved through *de-novo* synthesis of compatible solutes that are accumulated mainly in the cytosol and through accumulation of inorganic ions, which accumulate in the vacuole where larger changes in ionic strength can be tolerated. Compatible solutes, which are defined as small water-soluble molecules devoid of metabolic effects, can also function in osmoprotection of cellular structures, by interacting with membranes, protein complexes, or enzymes (Bohnert et al., 1995). OA, which occurred under all stress combinations, has increased with the aggravation of the stresses (Fig. 4A) and is proportional to the reduction in plant water status (Blum, 2017). In accordance, accumulation of compatible solutes such as glucose, proline and GABA, was positively correlated with OA, and exhibited a higher accumulation under S_&_D and S_&_D_&_H, and a lower accumulation under S_&_H and D_&_H (Fig. 4B and Supplementary Table S3). Plants’ ability to retain a high water content under S_&_H, despite a relative mild increase in organic osmolyte accumulation, may be attributed to an increase in inorganic osmolytes such as K^+^ and especially Na^+^ that are non-competitive to growth (Fig. 5C, and Supplementary Fig. S5). D*e novo* synthesis of osmolytes is an energetically expensive process that may come at a large metabolic cost. Therefore, ion accumulation can be an energetically more favorable option (providing it is effectively sequestered in the vacuole) (Fan et al., 2015). Accumulation of inorganic osmolytes may mitigate the energy trade-off between defense mechanisms and reproductive output and prioritize the latter, thus, minimizing yield penalty under S_&_H (Shabala and Shabala, 2011).

### Plants’ energy balance and fitness were severely damaged under combinations of S_&_D and S_&_D_&_H

Soil salinity is a growing concern in arid and semi-arid regions, both in cultivated land and natural habitats, leading to severe reductions in plant growth and productivity (Rozema and Flowers, 2008). Limited water availability can impair Hill reaction, evaporative coolant and mass flow of solvents, from both the soil and source tissues (Bohnert et al., 1995). Indeed, under S_&_D and S_&_D_&_H, plants were not able to retain a high RWC as control plants (Fig. 4B), and photosynthesis and transpiration rates dropped by ≥ 82 and 90%, respectively. In accordance, leaf temperature increased significantly compared with either control or single heat stress (Shaar-Moshe et al., 2017), implying limited ability of the plants to cool their leaves by transpiration. S_&_D and S_&_D_&_H also led to sever growth attenuations and to a decrease in the reproductive output by 63 and 74%, respectively (Fig. 1C-G). Similar effects of reduction in plant growth and productivity, were reported for wild (*Hordeum spontaneum*) and domesticated (*H. vulgare*) barley subjected to S_&_D during the vegetative and reproductive stages (Ahmed et al., 2013).

Sink strength is of a paramount importance for grain filling. The developing grains are mainly depended on two sources of assimilates: photosynthesis after anthesis and translocation of reservoirs assimilated before anthesis and temporarily stored in the stem. Under stress conditions, which diminish photosynthesis capacity of source tissues, grain filling is largely dependent on remobilization of stem reserves (Gallagher et al., 1976). The prolonged stress conditions, imposed under S_&_D and S_&_D_&_H, severely curtailed photosynthesis capacity and shifted the balance from maintenance and reproduction towards defense processes throughout the plants’ life-history. As a consequence of the decreased photosynthesis and culm length (Shaar-Moshe et al., 2017), these plants are expected to contain less stem reserves (Borrell et al., 1993) that are essential for grain filling and source-sink homeostasis (Jagadish et al., 2015).

The effect of phenotypic plasticity on plant fitness can be positive, neutral or maladaptive (van Kleunen and Fischer, 2005). Thus, not all alterations lead to the desired outcome of ameliorating the unfavorable conditions (Donohue, 2003). The limited water availability, under S_&_D and S_&_D_&_H, could result in attenuation of shoot DW accumulation (Fig. 1A), which leads to accumulation of sugars, such as glucose and galactose (Fig. 3 and Supplementary Table S3), due to a decreased demand (Hummel et al., 2010). Sugar accumulation can also result in feedback inhibition of photosynthesis (Rossi et al., 2015). Depletions of primary metabolites of the TCA cycle, such as citrate and succinate under S_&_D_&_H, and malate under S_&_D, further suggest that cellular respiration, were compromised under these combinations. GABA, which has been shown to accumulate under abiotic and biotic stresses and is associated with diverse physiological responses, including fluxes of carbon into the TCA cycle, nitrogen metabolism and osmoregulation (Bouche and Fromm, 2004), was largely produced under S_&_D and S_&_D_&_H (Fig. 3 and Supplementary Table S3). Since GABA shunt is closely connected to the TCA cycle (Fait et al., 2007), the surplus accumulation of GABA, under S_&_D and S_&_D_&_H, may result from a decline in respiration. Primary metabolism can also be modified due to increased activation of secondary metabolism under different environmental constrains (Kooke and Keurentjes, 2011).

Altogether, the morph-physiological traits and metabolic profiling demonstrate the aggravating effects of the combined stresses, which hamper plant performances, diminish their fitness and may impend their dispersion in areas prone to simultaneous occurrence of salinity and drought with or without heat stress.

### Conclusions and future perspective

The prevalence of different combinations of stresses in the field, and their projected increase in frequency and intensity (Suzuki et al., 2014; Zandalinas et al., 2018), challenges the common methodology of single stress treatments and the relevance of such assays to plant acclimation under natural and agricultural systems. The variable and unpredictable environmental conditions that are associated with climate change, force plants to rapidly acclimate, both within and across generations (Nicotra et al., 2010). Understanding the effects of environmental constraints on plant life-history strategies and the limits of plasticity under changing environments can assist in predicting species dispersion and facilitate the development of varieties, which are better adapted to the foreseen climatic conditions. To meet these challenges, a multidisciplinary approach that considers physiological and phenological information of realistic stress assays, together with system regulatory mechanisms, from the gene to the whole plant level, should be undertaken.

## Supplementary material

Supplementary data are available at JXB online.

**Table S1.**

Statistical analysis of physiological traits at three developmental stages.

**Table S2.**

List of differentially expressed genes that were used for correlation analyses.

**Table S3.**

List of fold change and false discovery rate values of putative metabolites.

**Fig. S1.**

A schematic overview of the stress combination assay.

**Fig. S2.**

Germination rates of seeds from plants subjected stress combinations.

**Fig. S3.**

Association between leaf starch content and physiological traits or gene expression pattern.

**Fig. S4.**

Galactose metabolic pathway under combinations of stresses.

**Fig. S5.**

Ion concentration in *Brachypodium distachyon* leaves under combinations of stresses.

## Acknowledgments

We would like to thank A. Oksenberg, I. Vilan, I. Ayalon and I. Sabag for their technical assistance with the physiological experiments and N. Jayasinghe for analysis of metabolic samples. GC-MS analysis was carried out at Metabolomics Australia, School of BioSciences, University of Melbourne, which is supported by funds from the Australian Government’s National Collaborative Research Infrastructure Scheme (NCRIS) administered through Bioplatforms Australia (BPA) Ltd. This research partially supported by the United States-Israel Binational Science Foundation (BSF) (grant #2011310) and the Hebrew University of Jerusalem. LSM is indebted to the Israeli President’s Scholarship for Scientific Excellence and Innovation.

## Author contributions

L.S.-M. and R.H. performed the experiments, L.S.-M. analyzed the experiments, L.S.-M., U.R., and Z.P. designed the experiments, L.S.-M., and Z.P. wrote the manuscript with edits from all the authors

## References

Ahmed IM, Cao F, Zhang M, Chen X, Zhang G, Wu F. 2013. Difference in yield and physiological features in response to drought and salinity combined stress during anthesis in Tibetan wild and cultivated barleys. PLoS One 8, https://doi.org/10.1371/journal.pone.0077869.

Avila-Ospina L, Moison M, Yoshimoto K, Masclaux-Daubresse C. 2014. Autophagy, plant senescence, and nutrient recycling. Journal of Experimental Botany 65, 3799-3811.

Barhoumi Z, Atia A, Rabhi M, Djebali W, Abdelly C, Smaoui A. 2010. Nitrogen and NaCl salinity effects on the growth and nutrient acquisition of the grasses *Aeluropus littoralis*, *Catapodium rigidum*, and *Brachypodium distachyum*. Journal of Plant Nutrition and Soil Science 173, 149-157.

Barry CS. 2009. The stay-green revolution: recent progress in deciphering the mechanisms of chlorophyll degradation in higher plants. Plant Science 176, 325-333.

Bazzaz FA, Chiariello NR, Coley PD, Pitelka LF. 1987. Allocating resources to reproduction and defense. Bioscience 37, 58-67.

Benjamini Y, Hochberg Y. 1995. Controlling the false discovery rate: A practical and powerful approach to multiple testing. Journal of the Royal Statistical Society. Series B (Methodological) 57, 289-300.

Blum A. 2017. Osmotic adjustment is a prime drought stress adaptive engine in support of plant production. Plant, Cell and Environment 40, 4-10.

Bohnert HJ, Nelson DE, Jensen RG. 1995. Adaptations to environmental stresses. Plant Cell 7, 1099-1111.

Borrell AK, Incoll LD, Dalling MJ. 1993. The influence of the *Rht1* and *Rht2* alleles on the deposition and use of stem reserves in wheat. Annals of Botany 71, 317-326.

Bouche N, Fromm H. 2004. GABA in plants: Just a metabolite? Trends in Plant Science 9, 110-115.

Bradshaw AD. 1965. Evolutionary significance of phenotypic plasticity in plants. Advances in Genetics 13, 115-155.

De Deyn GB. 2017. Plant life history and above-belowground interactions: missing links. Oikos 126, 497-507.

Des Marais DL, Juenger TE. 2016. *Brachypodium* and the abiotic environment. In: Vogel J, ed. Genetics and Genomics of Brachypodium. Switzerland: Springer, 291-311.

Des Marais DL, Lasky JR, Verslues PE, Chang TZ, Juenger TE. 2017. Interactive effects of water limitation and elevated temperature on the physiology, development and fitness of diverse accessions of *Brachypodium distachyon*. New Phytologist 214, 132-144.

Dias DA, Hill CB, Jayasinghe NS, Atieno J, Sutton T, Roessner U. 2015. Quantitative profiling of polar primary metabolites of two chickpea cultivars with contrasting responses to salinity. Journal of Chromatography B 1000, 1-13.

Donohue K. 2003. Setting the stage: phenotypic plasticity as habitat selection. International Journal of Plant Sciences 164, S79-S92.

Fait A, Fromm H, Walter D, Galili G, Fernie AR. 2007. Highway or byway: the metabolic role of the GABA shunt in plants. Trends in Plant Science 13, 14-19.

Fan Y, Shabala S, Ma Y, Xu R, Zhou M. 2015. Using QTL mapping to investigate the relationships between abiotic stress tolerance (drought and salinity) and agronomic and physiological traits. BMC Genomics 16, 43.

Gallagher JN, Biscoe PV, Hunter B. 1976. Effects of drought on grain-growth. Nature 264, 541-542.

Gitelson AA, Zur Y, Chivkunova OB, Merzlyak MN. 2002. Assessing carotenoid content in plant leaves with reflectance spectroscopy. Photochemistry and Photobiology 75, 272-281.

Grime JP. 1977. Evidence for the existence of three primary strategies in plants and its relevance to ecological and evolutionary theory. American Naturalist 111, 1169-1194.

Hill CB, Dias DA, Roessner U. 2015. Current and emerging applications of metabolomics in the field of agricultural biotechnology. In: Ravishankar RV, ed. Advances in Food Biotechnolog. John Wiley & Sons Ltd, 13–26.

Hirschberg J. 2001. Carotenoid biosynthesis in flowering plants. Current Opinion in Plant Biology 4, 210-218.

Holzl G, Dormann P. 2007. Structure and function of glycoglycerolipids in plants and bacteria. Progress in Lipid Research 46, 225-243.

Hong SY, Park JH, Cho SH, Yang MS, Park CM. 2011. Phenological growth stages of *Brachypodium distachyon*: codification and description. Weed Research 51, 612-620.

Hummel I, Pantin F, Sulpice R, et al.. 2010. *Arabidopsis* plants acclimate to water deficit at low cost through changes of carbon usage: an integrated perspective using growth, metabolite, enzyme, and gene expression analysis. Plant Physiology 154, 357-372.

Jagadish KSV, Kishor PBK, Bahuguna RN, von Wiren N. Sreenivasulu N. 2015. Staying alive or going to die during terminal senescence-an enigma surrounding yield stability. Frontiers in Plant Science 6, 1070.

Jongebloed U, Szederkenyi J, Hartig K, Schobert C, Komor E. 2004. Sequence of morphological and physiological events during natural ageing and senescence of a castor bean leaf: sieve tube occlusion and carbohydrate back-up precede chlorophyll degradation. Physiologia Plantarum 120, 338-346.

Jump AS, Peñuelas J. 2005. Running to stand still: adaptation and the response of plants to rapid climate change. Ecology Letters 8, 1010-1020.

Kooke R, Keurentjes JJB. 2011. Multi-dimensional regulation of metabolic networks shaping plant development and performance. Journal of Experimental Botany 63, 3353-3365.

Krasensky J, Jonak C. 2012. Drought, salt, and temperature stress-induced metabolic rearrangements and regulatory networks. Journal of Experimental Botany 63, 1593–1608

Levitt J. 1972. Responses of Plant to Environmental Stress. N.Y. Academic Press.

Loss SP, Siddique KHM. 1994. Morphological and physiological traits associated with wheat yield increases in Mediterranean environments. Advances in Agronomy 52, 229-276.

Love MI, Huber W, Anders S. 2014. Moderated estimation of fold change and dispersion for RNA-seq data with DESeq2. Genome Biology 15, 550.

Lu Y. 2016. Identification and roles of photosystem II assembly, stability, and repair factors in Arabidopsis. Frontiers in Plant Science 7,168.

Maggio A, De Pascale S, Fagnano M, Barbieri G. 2011. Saline agriculture in Mediterranean environments. Italian Journal of Agronomy 6, 36-43.

Matesanz S, Gianoli E, Valladares F. 2010. Global change and the evolution of phenotypic plasticity in plants. Annals of the New York Academy of Sciences 1206, 35-55.

Merzlyak MN, Gitelson AA, Chivkunova OB, Rakitin VY. 1999. Non destructive optical detection of pigment changes during leaf senescence and fruit ripening. Physiologia Plantarum 106, 135-141.

Mickelbart MV, Hasegawa PM, Bailey-Serres J. 2015. Genetic mechanisms of abiotic stress tolerance that translate to crop yield stability. Nature Reviews Genetics 16, 237-251.

Naschitz S, Naor A, Genish S, Wolf S, Goldschmidt EE. 2010. Internal management of non-structural carbohydrate resources in apple leaves and branch wood under a broad range of sink and source manipulations. Tree Physiology 30, 715-727.

Nicotra AB, Atkin OK, Bonser SP, Davidson AM, Finnegan EJ, Mathesius U, van Kleunen M. 2010. Plant phenotypic plasticity in a changing climate. Trends in Plant Science 15, 684-692.

Nilson SE, Assmann SM. 2010. Heterotrimeric G proteins regulate reproductive trait plasticity in response to water availability. New Phytologist 185, 734-746.

Nisar N, Li L, Lu S, Khin NC, Pogson BJ. 2015. Carotenoid metabolism in plants. Molecular Plant 8, 68-82.

Pigliucci M. 2001. Phenotypic Plasticity: Beyond Nature and Nurture. Baltimore, MD: The Johns Hopkins University Press.

Rontein D, Basset G, Hanson AD. 2002. Metabolic engineering of osmoprotectant accumulation in plants. Metabolic Engineering 4, 49-56.

Rossi M, Bermudez L, Carrari F. 2015. Crop yield: challenges from a metabolic perspective. Current Opinion in Plant Biology 25, 79-89.

Rozema J, Flowers T. 2008. Crops for a salinized world. Science 322, 1478-1480.

Ruan Y.L 2014. Sucrose metabolism: gateway to diverse carbon use and sugar signaling. Annual Review of Plant Biology 65, 33-67.

Sade N, Del Mar Rubio-Wilhelmi M, Umnajkitikorn K, Blumwald E. 2018. Stress-induced senescence and plant tolerance to abiotic stress. Journal of Experimental Botany 69, 845–853.

Salguero-Gomez R, Jones OR, Jongejans E, Blomberg SP, Hodgson DJ, Mbeau-Ache C, Zuidema PA, de Kroon H, Buckley YM. 2016. Fast-slow continuum and reproductive strategies structure plant life-history variation worldwide. Proceedings of the National Academy of Sciences, USA 113, 230-235.

Schlichting CD, Smith H. 2002. Phenotypic plasticity: Linking molecular mechanisms with evolutionary outcomes. Evolutionary Ecology 16, 189-211.

Scholthof KBG, Irigoyen S, Catalan P, Mandadi K. 2018. *Brachypodium*: A monocot grass model system for plant biology. Plant Cell, 30, 1673-1694.

Shaar-Moshe L, Blumwald E, Peleg Z. 2017. Unique physiological and transcriptional shifts under combinations of salinity, drought, and heat. Plant Physiology 174, 421-434.

Shaar-Moshe L, Hubner S, Peleg Z. 2015. Identification of conserved drought-adaptive genes using a cross-species meta-analysis approach. BMC Plant Biology 15, 111.

Shabala S, Shabala L. 2011. Ion transport and osmotic adjustment in plants and bacteria. Biomolecular Concepts 2, 407-419.

Sultan SE. 2000. Phenotypic plasticity for plant development, function and life history. Trends in Plant Sciences 5, 537-542.

Suzuki N, Rivero RM, Shulaev V, Blumwald E, Mittler R. 2014. Abiotic and biotic stress combinations. New Phytologist 203, 32-43.

Thalmann M, Pazmino D, Seung D, Horrer D, Nigro A, Meier T, Kölling K, Pfeifhofer HW, Zeeman SC, Santelia D. 2016. Regulation of leaf starch degradation by abscisic acid is important for osmotic stress tolerance in plants. Plant Cell 28, 1860-1878.

Thalmann M, Santelia D. 2017. Starch as a determinant of plant fitness under abiotic stress. New Phytologist 214, 943-951.

Thimm O, Blasing O, Gibon Y, Nagel A, Meyer S, Kruger P, Selbig J, Muller LA, Rhee SY, Stitt M. 2004. MAPMAN: A user-driven tool to display genomics data sets onto diagrams of metabolic pathways and other biological processes. Plant Journal 37, 914-939.

van Kleunen M, Fischer M. 2005. Constraints on the evolution of adaptive phenotypic plasticity in plants. New Phytologist 166, 49-60.

Vautard R, Yiou P, D’Andrea F, de Noblet N, Viovy N, Cassou C, Polcher J, Ciais P, Kageyama M, Fan Y. 2007. Summertime European heat and drought waves induced by wintertime Mediterranean rainfall deficit. Geophysical Research Letters 34, L07711

Violle C, Navas ML, Vile D, Kazakou E, Fortunel C. Hummel I, Garnier E. 2007. Let the concept of trait be functional! Oikos 116, 882-892.

Watt M, Moosavi S, Cunningham SC, Kirkegaard JA, Rebetzke GJ, Richards RA. 2013. A rapid, controlled-environment seedling root screen for wheat correlates well with rooting depths at vegetative, but not reproductive, stages at two field sites. Annals of Botany 112, 447-455.

Whan AP, Smith AB, Cavanagh CR, Ral JP, Shaw LM, Howitt CA, Bischof L. 2014. GrainScan: A low cost, fast method for grain size and colour measurements. Plant Methods 10, 23.

Xia J, Wishart DS. 2016. Using MetaboAnalyst 3.0 for comprehensive metabolomics data analysis. Current Protocols in Bioinformatics 55, 10-14.

Yermiyahu U, Heuer B, Silverman D, Faingold I, Avraham L. 2017. Nitrate analysis of *Diplotaxis tenuifolia*: Fresh versus dry material for meeting international standards and regulations. Israel Journal of Plant Sciences, 19, 1-4.

Zandalinas SI, Mittler R, Balfagon D, Arbona V, Gomez-Cadenas A. 2018. Plant adaptations to the combination of drought and high temperatures. Physiologia Plantarum 162, 2-12.

